# Deletions of distant regulatory sequences upstream of zebrafish *pitx2* result in a range of ocular phenotypes

**DOI:** 10.1101/772426

**Authors:** Eric Weh, Elena Sorokina, Kathryn Hendee, Doug B. Gould, Elena V. Semina

**Author notes:** first co-authors (Eric Weh, Elena Sorokina).

## Abstract

Development of the anterior segment of the vertebrate eye is a highly coordinated process. Genetic mutations in factors guiding this process result in Anterior Segment Dysgenesis (ASD), a spectrum of disorders affecting the iris, cornea, trabecular meshwork and/or other iridocorneal angle structures and associated with glaucoma. One of the first factors linked to ASD in humans was *PITX2*, a homeodomain containing transcription factor with a role in Axenfeld-Rieger syndrome (ARS). In addition to pathogenic alleles within the coding region of *PITX2*, deletions affecting the distant upstream region, but not *PITX2* itself, have also been reported in ARS. Consistent with this, the distant upstream region was shown to contain multiple conserved elements (CE) with *pitx2*-related enhancer activity identified through studies in zebrafish. The two smallest human deletions reported to date encompass conserved elements 5-11 (*ΔCE5-11*) or 5-7 (*ΔCE5-7*). We previously reported the generation of *ΔCE5-11* in zebrafish and we have now replicated the smallest deletion, *ΔCE5-7*, in the same model and studied the associated phenotype, expression, and DNA methylation profiles; we also performed further phenotypic examinations of the *pitx2^ΔCE5-11^* fish. We show that the expression changes and phenotypes observed in the two lines are variable but that the severity generally correlates with the size of the deletion and the number of affected CEs; *pitx2* promoter and a nearby region were hypermethylated in the *pitx2^ΔCE5-7^* embryonic eyes. In addition, a subset of *pitx2^ΔCE5-11^* animals were found to have a severe retinal phenotype suggesting that additional factors may modify the effects of this allele. These data provide further insight into functional sequences in the *PITX2/pitx2* genomic region that coordinate *PITX2/pitx2* expression during eye development and provide the basis for future studies into *PITX2/pitx2* upstream regulators and modifiers.

## INTRODUCTION

Axenfeld-Rieger Syndrome (ARS) is an autosomal dominant disorder characterized by Anterior Segment Dysgenesis (ASD), dental anomalies and periumbilical skin defects. Mutations within *PITX2* and several other factors have been reported as causative in patients with ARS and other ASD conditions. The spectrum of ASD in individuals with *PITX2* mutations ranges from mild (isolated posterior embryotoxon) to severe (aniridia) with a typical presentation consisting of iris hypoplasia, posterior embryotoxon, and irido-corneal adhesions; developmental glaucoma is strongly associated with all of these phenotypes (Reis and Semina, 2011; Reis et al. 2012; Hendee et al. 2018).

The development of the anterior chamber begins shortly after separation of the lens vesicle from the surface ectoderm. At this time the cells surrounding the optic cup, termed the periocular mesenchyme, begin migrating in towards the space between the lens and what will become the corneal epithelium. These cells migrate in waves, number of which vary with the particular species, to create the corneal endothelium and stroma followed by the iris stroma. In mice, the major structures of the anterior chamber are present by e15.5 with the irido-corneal angle fully formed by approximately p21. The major morphogenic processes are similar in zebrafish. The structures of the anterior segment are formed by approximately 5 days post fertilization (dpf) with morphogenesis proceeding over the first month of life (Cvekl and Tamm, 2004; Soules and Link, 2005). Consistent with its role in ASD in humans, the transcription factor *pitx2/pitx2* was shown to serve as a key effector during these developmental processes (Hjalt et al., 2000; Sowden, 2007; Volkmann et al., 2011; Liu and Semina 2012; Chen and Gage 2016; Hendee et al. 2018). *PITX2/pitx2* expression begins early in the migrating periocular mesenchyme cells, soon after the separation of the lens vesicle from the surface ectoderm (e10.5 in mice, ~22-hpf in zebrafish). As ocular development proceeds, *pitx2/pitx2* expression becomes more restricted to the cornea, iris and irido-comeal angle structures.

The exact transcriptional processes which coordinate the expression of *pitx2/pitx2* during development remain largely unknown. Previous studies have demonstrated the importance of retinoic acid (RA) signaling for neural crest cells during ocular development (Gage and Zacharias, 2009; Matt et al., 2008). *Pitx2* is expressed in the periocular mesenchyme, which is comprised of neural crest and mesodermal cells. Matt and colleagues found that inactivating all three retinoic acid receptors (RARs) specifically in neural crest cells leads to a loss of *Pitx2* expression within the periocular mesenchyme but not in the developing extraocular muscles. These data demonstrate the importance of RA signaling for the initiation of *Pitx2* expression; however, it is unclear whether RA signaling is sufficient for *Pitx2* expression or if other factors are involved in controlling *Pitx2* transcription during ocular development. Additional studies have shown that normal WNT/β-catenin signaling is essential for the maintenance of *Pitx2* expression during later stages of ocular development (Zacharias and Gage, 2010). Similar to Matt and colleagues, Zacharias and Gage inactivated β-catenin specifically in neural crest cells. They found that *Pitx2* is expressed at e10.5 but is then quickly lost as development continues, suggesting that WNT/β-catenin signaling is essential for the maintenance of *Pitx2* expression after activation by RA signaling. Interestingly, two WNT antagonists, *Dkk2* and *notum*, were shown to be downstream of Pitx2/pitx2 with *Dkk2* demonstrated to be a direct target (Gage et al. 2008; Hendee et al. 2019). Currently available data are not sufficient to fully understand the transcriptional landscape governing the spatio-temporal expression pattern of *pitx2/pitx2*, therefore additional studies are necessary to understand what regulates its expression.

Previous studies identified multiple conserved DNA elements (CEs) upstream of *PITX2/pitx2* and additional CEs within introns; each of these elements was preserved in the genomes of numerous species studied in both their number and organization (Volkmann et al. 2011). *In vivo* reporter expression found that many of these CEs are capable of driving GFP in a pattern consistent with that of endogenous *pitx2*, including ocular expression (Volkmann et al., 2011). Additional evidence from human patients with ARS demonstrated that deletions of these upstream regulatory elements are sufficient to produce the ARS phenotype (Ansari et al., 2016; Reis et al., 2012; Volkmann et al., 2011; Protas et al. 2017). The two smallest overlapping deletions which affect from three to seven of these elements, CE5-7 and CE5-11, were reported by Protas et al.; the zebrafish model was utilized to replicate the CE5-11 deletion and demonstrated a reduction of the anterior segment space in the mutant animals indicative of a developmental defect (Protas et al. 2017).

Here, we report the generation of a zebrafish line carrying a deletion of the upstream *pitx2* region comprising CE5-7 and thus replicating the smallest human deletion reported to date (Protas et al. 2017). We show that deletion of CE5-7 results in significant changes in *pitx2* expression associated with hypermethylation at the *pitx2* promoter/regulatory region and resulting in a reduction of anterior segment space in embryos and defects in the development of the iridocorneal angles identified by electron microscopy. Additionally, we performed further studies of the previously reported zebrafish line carrying the larger CE5-11 deletion and show that the expression changes and phenotypes observed in the two regulatory lines show variable severity correlating with the number of affected CEs. Also, a subset of *pitx2^ΔCE5-11^* animals with a more severe retinal phenotype was identified suggesting a sensitizing effect of incomplete *pitx2* deficiency and a role of additional modifiers in its phenotypic expression. In summary, the presented data provide further insight into the regulatory elements of *PITX2/pitx2* and highlight their conserved function. Identifying the factors which bind to these elements, as well as modifiers of *pitx2*-associated phenotypes, will inform our understanding of ocular development and potentially reveal novel genes implicated in ASD.

## METHODS

### Animals

All animal studies were conducted with approval from the animal care and use committee at the Medical College of Wisconsin. Zebrafish, *Danio rerio*, were maintained on a 14/10 hour light/dark cycle. Embryos were raised at 28.5°C in standard E2 medium. When necessary, embryos were grown in E2 medium containing 0.003% phenylthiourea (PTU) to maintain transparency during the first 7 days of development.

### Zebrafish Line Generation

sgRNA was designed using the ZiFiT targeting software (http://zifit.partners.org/ZiFiT/) (Sander et al., 2007; Sander et al., 2010) using the T7 promoter settings. sgRNA (Table S1) were chosen which flank the desired conserved element for removal. Oligos were ordered exactly as displayed by the ZiFiT software from Eurofins MWG Operon (Thermo Fisher Scientific, Waltham, MA, USA). Oligos were annealed and ligated into the DR274 vector, a gift from Keith Joung (Addgene plasmid # 42250), following the protocol outlined by Hwang et al. (Hwang et al., 2013). Ligated plasmids were then cloned into DH5α Max Efficiency cells (Thermo Fisher Scientific, Waltham, MA, USA). Two colonies were picked for each plasmid, grown overnight and plasmid DNA was extracted using the PureLink Quick Plasmid Miniprep Kit (Thermo Fisher Scientific, Waltham, MA, USA). Plasmids were Sanger sequenced using the M13F sequencing primer and correct sgRNA sequences were identified using a BLAST alignment of the sgRNA oligos. Correct plasmids were then digested using Dral (New England Biolabs, Ipswich, MA, USA). Linearized plasmid was used as a template for RNA synthesis using the MEGAshortscript T7 Transcription Kit following the manufacturer's instructions (Thermo Fisher Scientific, Waltham, MA, USA). sgRNA was purified using the RNA Clean and Concentrator Kit (Zymo Research, Irvine, CA, USA) after DNA digestion using DNAse I, amplification grade (Thermo Fisher Scientific, Waltham, MA, USA). mRNA for Sp.Cas9 was synthesized using linearized MLM3639, a gift from Keith Joung (Addgene plasmid # 42252), following the protocol outlined by Hwang et al. (Hwang et al., 2013) using the mMessage mMachine T7 Transcription Kit following the manufacturer's instructions (Thermo Fisher Scientific, Waltham, MA, USA). mRNA was modified to contain a poly-A tail using the Poly(A) Tailing Kit (Thermo Fisher Scientific, Waltham, MA, USA) following the manufacturer's instructions. Finally, poly-adenylated Cas9 mRNA was purified using the RNA Clean and Concentrator Kit (Zymo Research, Irvine, CA, USA).

Injections into zebrafish embryos were performed using a Nanoject II (Drummond Scientific, Broomall, PA, USA) set to deliver 9.2nL of injection solution. Injection solution was buffered using PBS and dyed with Phenol Red to visualize injections. The final injection concentrations were as follows: 10.8ng/μL each of sgRNA, 108ng/μL Cas9 mRNA, 54.4ng/μL of eGFP-Nanos3 mRNA. mRNA generated from a plasmid encoding for eGFP containing the 3'UTR of the Nanos3 gene was used to screen for zebrafish where injected mRNA was delivered to primordial germ cells to enhance germline transmission rates (Dong et al., 2014). Briefly, embryos injected with sgRNA/Cas9/eGFP-Nanos3 were screened at 48-hpf for GFP fluorescence in primordial germ cells. Embryos positive for staining were selected and raised to maturity. Embryos which were negative for staining were euthanized. Approximately 20 embryos from each injection were screened via PCR to detect the presence of large deletions induced by CRISPR editing events. Genomic DNA was extracted from embryos at 48-hpf using alkaline lysis as described previously (Meeker et al., 2007). PCR primers (Table S1) flanking the sgRNA sites were used to amplify a product only when a deletion removed the intervening DNA. Amplified products were Sanger sequenced to confirm deletion of the desired conserved elements. Presumed mosaic adults were in-crossed in pairs and approximately 24 embryos from each pair were screened for a deletion as described above and a mosaic adult for CE5-7 deletion was identified. The mosaic adult was outcrossed to produce embryos that were raised to maturity, genotyped to identify deletion carriers and in-crossed to generate homozygous embryos for further studies. The CE5-11 deletion line has been published previously (Protas et al. 2017).

### Light Microscopy and histology

*In vivo* imaging was performed on 1xTricane anesthetized embryos embedded in 1% low-melting point agarose. Adult fish were also anesthetized by 1xTricane in Ringer buffer and positioned on the plate filled with agarose gel. Imaging was done with either Stereo Discovery.V12microscope (Zeiss) for black and white images or Nikon SMZ1500 microscope (Nikon, Japan) equipped with an HR Plan Apo 1x WD54 objective for color images. For histology, embryos or adult fish were fixed for 1-2 days in modified Davidson solution (glacial acetic acid, 1 part, 95% ethyl alcohol, 1 part, 10% neutral buffered formalin, 2 parts, distilled water, 3 parts), rinsed in PBS and placed in 70% ethanol. Paraffin blocks were prepared, sectioned at 4μn and stained with hematoxylin and eosin in the Children's Research Institute Histology Core at the Medical College of Wisconsin. Slides were scanned on a NanoZoomer digital slide scanner (Hamamatsu, Hamamatsu City, Japan).

### Anterior chamber of the eye measurement

Embryos from heterozygous in-crosses of either the *pitx2^ΔCE5-7^* or *pitx2^ΔCE5-11^* lines were anesthetized using Tricane (MS-222). For anterior chamber measurements, embryos were individually anesthetized immediately before mounting in 1% low melt agar. The anterior chamber of embryos was measured using ImageJ as previously described (Protas et al. 2017). Briefly, we outlined the interior surface of the cornea, lens and iris to obtain the area measurement. We then measured the width of the eye at the widest point. Each measurement was made three times and the average was taken. The average anterior chamber area was divided by the average width of the eye for normalization. In order to eliminate bias in measurements, the genotype of each embryo was not determined until after all images were taken and measurements completed. Embryos were genotyped as outlined above and measurements were then binned to their respective genotype. Significance was determined using a two-tailed, un-equal variance student's t-test in Excel.

### qRT-PCR Analysis

The relative abundance of the two splice isoforms of *pitx2* (*pitx2a* and *pitx2c*) were measured using PowerUp Sybr Green Master Mix (Thermo Fisher Scientific, Waltham, MA, USA) and a BioRad CFX 96 or 384 Real-Time PCR Detection System (Bio-Rad). The primers and methods used to measure *pitx2a* and *pitx2c* levels, as well as *β-actin* (control) were described previously (Protas et al., 2017). Homozygous deletion (both lines) and wildtype embryos were dissected to collect eyes only for RNA extraction, while the remaining tissues were used for genotyping. Following genotyping, total RNA was extracted from at least 30 eyes for each developmental stage point/genotype. Tissues were solubilized in TRI Regent (Zymo Research, Irvine, CA, USA) and RNA was extracted using the Direct-Zol RNA Extraction Kit (Zymo research, Irvine, CA, USA) following the manufacturer's instructions. Purified total RNA samples were treated with DNAse I, Amplification Grade (Thermo Fisher Scientific, Waltham, MA, USA) following the manufacturer's instructions. DNAse treated total RNA was then purified using the RNA Clean and Concentrator Kit (Zymo Research, Irvine, CA, USA) and finally quantitated using a NanoDrop 1000 (Thermo Fisher Scientific, Waltham, MA, USA). 150 ng of total RNA was used as template for reverse transcription using the superScript III First-Strand Synthesis System (Thermo Fisher Scientific, Waltham, MA, USA) following the manufacturer's instructions, β-actin was used to normalize cycle values. Expression levels in wild-type tissues were used as the reference in comparisons and data were compared using a two-tailed, unpaired student's t-test.

### Electron microscopy

Sections of the ventral and dorsal angle of the zebrafish eye as well as whole corneas were examined by electron microscopy at 3-dpf and 14-dpf in homozygous *pitx2^ΔCE5-7^* and *pitx2^ΔCE5-11^* as well as WT eyes. Embryos were fixed for 24 hours at 4°C in primary fixative (2% paraformaldehyde, 2.5% glutaraldehyde, 3% sucrose, 0.06% phosphate buffer, pH 7.4), washed and post-fixed with 1% osmium tetroxide on ice as described in Hendee et al, 2018. After dehydration by submerging the embryos in a series of the methanol and acetonitrile solutions, specimens were infiltrated with the EMbed-812resin mixture (14120; Electron Microscopy Sciences, Hatfield, PA). Semi-thin transverse sections (500 nm) were cut throughout the eye and stained with 1% Toluidine Blue in 1% Borax buffer until reaching the center of the lens. Ultra-thin sections (70–80 nm) were collected on copper hexagonal mesh coated grids (G200H-Cu; Electron Microscopy Sciences). Grids were stained with uranyl acetate and lead citrate. Images were taken using a Hitachi H600 TEM microscope (Hitachi, Tokyo, Japan).

### Whole Genome Bisulfite Sequencing (WGBS)

25-30 eyes were dissected from both WT and *pitx2^ΔCE5-7^* eye zebrafish embryos at 3 dpf in Ringer buffer and genomic DNA was purified using Quick-DNA microprep Plus kit (ZYMO Research). Eyes were resuspended in Solid Tissue buffer containing Proteinase K and incubated at 55°C for 6 hours. Two volumes of the Genomic Binding buffer were added to the solution and applied to the Zymo-Spin IC-XM column. The further steps were performed following manufacturer protocol.

For WGBS analysis DNA was submitted to the Macrogen NGS service (Macrogen, Rockville). After converting unmethylated cytosines to uracils (EZ DNA Methylation-Gold™ Kit (Zymo Research)), indexed libraries were prepared using Accel-NGS Methyl-Seq DNA Library Kit EZ DNA (Swift Biosciences, Inc.) as outlined in the manufacturers’ protocol. Briefly, genomic DNA samples from 3 biological replicates for both WT and *pitx2^ΔCE5-7^ eye* RNA samples were subjected to bisulfite-mediated cytosine to uracil conversion. Bisulfite conversion rate (%) for all samples was 99.7%. DNA fragments were treated with adaptase to achieve simultaneous tailing and ligation of truncated adapter 1 to the 3’ ends of DNA fragments. Then, extension and ligation of the second truncated adaptor 2 to the bottom strand was performed and the library was amplified and indexed. The high throughput sequencing was performed on the lllumina platform with average read length of 151 bases. Reads were truncated in order to eliminate adapter sequences and bases with low quality using the Trim Galore 0.4.5 program. Only reads longer then 20bp were selected for further analysis. In all samples 97% of the reads had phred quality score greater than or equal to 20 as defined by FastQC v0.11.5 program. The trimmed reads were mapped to the zebrafish GRCz11 reference genome assembly with BSMAP v2.87 which is based on SOAP (ShortOligo Alignment Program). Index sequences and PCR duplicateswere removed with SAMBAMBA (v0.5.9). The methylation ratio for each cytosine in the CG (or CpG), CHG or CHH sites was defined using 'methylatio.py’ script in BSMAP. These values were normalized using median scaling normalization and statistical analysis of the pairs was performed. The significant results were selected on the conditions of | delta_mean | >=0.2 & raw p-value<0.05.

## RESULTS

### Generation of a zebrafish line carrying a CE5-7 deletion of pitx2 upstream region

We targeted the upstream region of *pitx2* containing conserved elements (CE) 5, 6 and 7 (CE5-7) for deletion using the CRISPR/Cas9 system (Figure 1). The CE5 and CE7 elements were previously shown to play a role in regulation of *pitx2* expression during eye development in zebrafish and a deletion involving these elements was reported in a patient affected with ARS (Volkmann et al. 2011; Protas et al. 2017). Founder fish carrying the CE5-7 deletion were identified using PCR and subsequent sequencing determined that the deletion was ~61 kb spanning chr14:36087688-36148476 (Figure 1) and the line was named *pitx2^ΔCE5-7^*. Heterozygous *pitx2^ΔCE5-7^* adult fish were crossed to generate animals homozygous for the deletion; homozygous embryos were present in expected Mendelian ratios (25%; n=64) and had no gross morphological defects, similar to the previously reported *pitx2^ΔCE5-11^* larvae with a larger regulatory deletion (Figure 2A; Protas et al. 2017). The *pitx2^ΔCE5-11^* animals carry a larger deletion of ~132 kb (chr14: 36014929-36146797) disrupting CE5-11 upstream of *pitx2* (Protas et al. 2017). Measurements of the anterior segment were performed on embryos produced by heterozygous *pitx2^ΔCE5-7^* parents and included a mix of siblings heterozygous or homozygous for the deletion allele as well as wild-type (WT) at 3-, 4-, and 5-dpf. The imaging and measurements (Figure 2B) were performed in a blind manner (prior to genotyping); following this, embryos were genotyped, grouped accordingly, and statistical analysis was performed. This analysis identified statistically significant differences between wild-type and homozygous eyes at 3-dpf (p=0.0019), 4 dpf (p=0.000044) and 5 (p=0.000051) and no statistically significant difference between wild-type and heterozygotes eyes (3 dpf: p=0.019; 4 dpf: p=0.898; 5dpf: p=0.985) (Figure 2C). Since heterozygote embryos showed no phenotype, we focused on homozygous embryos for further analysis. A similar, though more significant, reduction in the anterior segment space was previously reported in other *pitx2* lines, including the loss-of-function allele *pitx2^M64*^* and regulatory deletion *pitx2^ΔCE5-11^* (Hendee et al. 2018, Protas et al. 2017) (see *pitx2^ΔCE5-7^* and *pitx2^ΔCE5-11^* mutants as well as WT side by side for comparison in Figure 2D-I) and suggests abnormal development of the anterior segment structures.

**Figure 1.**
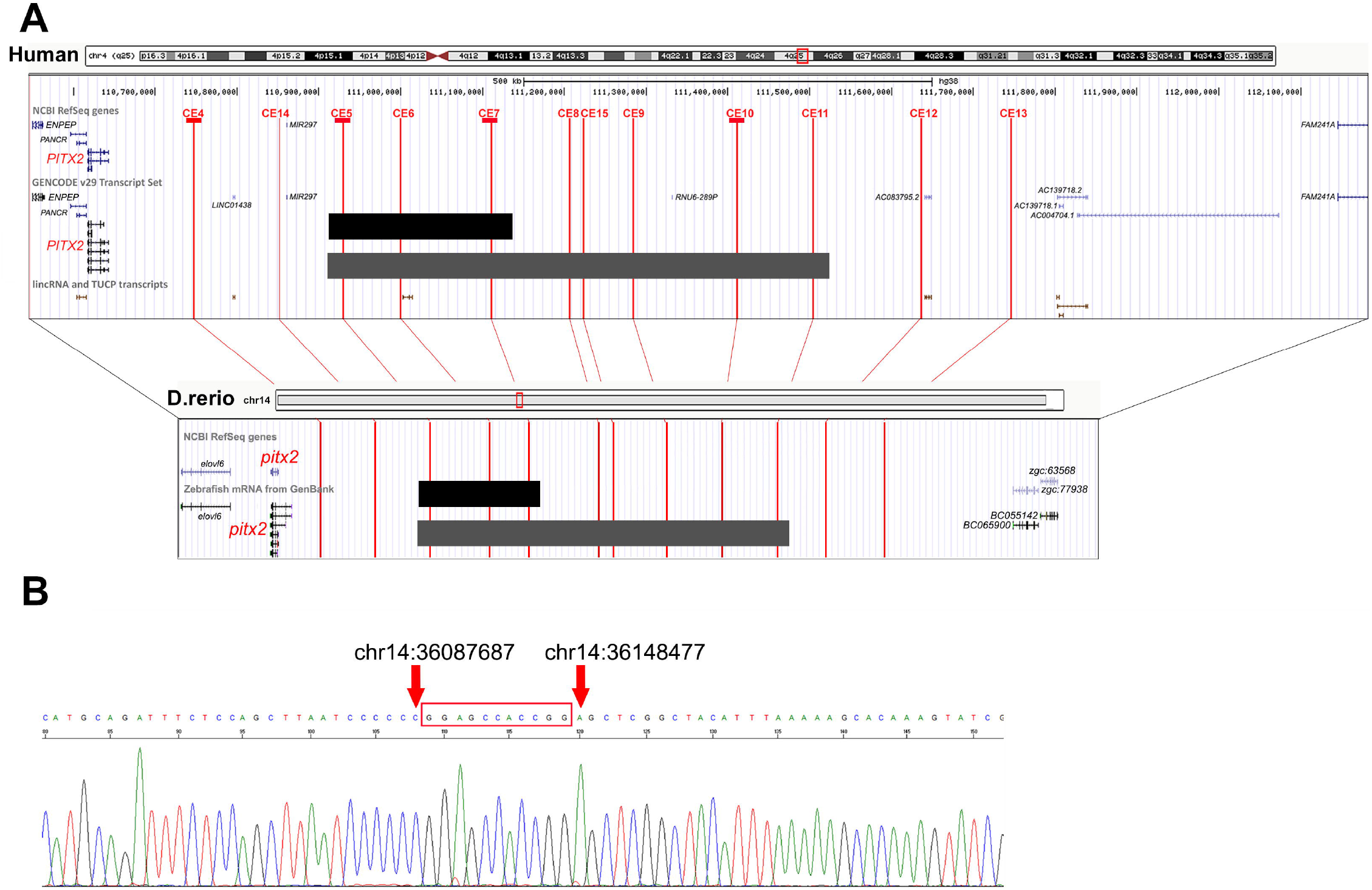
Schematic drawing of the human 4q25 region and the orthologous zebrafish chromosome 14 region. **(A)** Genome Browser (https://genome.ucsc.edu/) views of the human 4q25 region encompassing *PITX2* and the nearest protein-encoding transcripts on each side, as well as the Danio rerio (zebrafish) chromosome 14 region containing *pitx2* and the adjacent protein-encoding transcripts on both sides. Please note the gene desert located in the 5'region of both human and zebrafish *PITX2/pitx2* genes that contains multiple distant conserved elements (CE) (Volkmann et al. 2012; Protas et al. 2017) marked with red lines and numbered CE4-CE15; CE4, 5, 7 and 10 associated with ocular expression are underscored. CE5-7 and CE5-11 deletions in human patients and zebrafish lines, presented previously (Protas et al. 2017) and in this report are shown as black and grey bars, respectively. Human *PITX2* is bordered by *PANCR* (PITX2 adjacent non-coding RNA) and *ENPEP* (glutamyl aminopeptidase) on its 3’ end (at a distance of ~2 kb and ~54 kb respectively) and *FAM241A* (family with sequence similarity 241 member A) on its 5’ end (~1503 kb from *PITX2).* The 5’ region contains *MIR297* microRNA located ~ 219 kb upstream along with several additional transcripts like *LINC01438* (long intergenic non-protein coding RNA 1438), *RNU6-289P* (RNA, U6 small nuclear 289, pseudogene), Gencode transcripts *AC083795.2, AC139718.2, AC139718.1, AC004704.1*, and several lincRNAs (large intergenic noncoding RNAs) and TUCPs (transcripts of uncertain coding potential). In zebrafish, *pitx2* is surrounded by *zgc:77938* (predicted to encode a gene orthologous to human *ADH4* (alcohol dehydrogenase 4 (class II), pi polypeptide)) at its 5’ end at a distance of 309 bp and *elovl6* (encodes ELOVL fatty acid elongase 6) at its 3’ end at distance 17 kb (human *ELOVL6* and *ADH4* are located at 418 kb and 11.5 Mb 3’ of PITX2). All data are presented according to Zebrafish May 2017 (GRCz11/danRer11) and Human Dec. 2013 (GRCh38/hg38) assemblies. **(B)** DNA sequence across the CE5-7 deleted region in *pitx2^ΔC5-7^* homozygous animals; genomic coordinates in the wild-type allele are indicated: in wild-type animals the region between the arrows spans 60,790 bp encompassing CEs 5, 6 and 7. This 60,790 bp sequence is deleted in the mutant allele and replaced with a 11-bp insertion (red box).

### Expression analysis of *pitx2* transcripts in mutant lines

Zebrafish *pitx2* encodes two isoforms, *pitx2a* and *pitx2c*, that share the same homeodomain and C-terminal sequence but have dissimilar N-terminal parts; both transcripts are active during ocular development (Liu and Semina 2012). To investigate the effect of the upstream deletion on *pitx2* expression, we tested transcript levels of both the *pitx2a* and *pitx2c* isoforms in *pitx2^ΔCE5-7^* and WT embryonic eyes via qPCR; independent ocular samples from the previously reported *pitx2^ΔCE5-11^* line served as a positive control. These experiments identified a statistically significant decrease in both *pitx2a* and *pitx2c* transcript levels in *pitx2^ΔCE5-7^* embryonic eyes at 4-dpf in comparison to WT levels; At 2- and 3-dpf, some changes in expression were also noted but these findings were not statistically significant (Figure 2J). Similar to the anterior segment reduction, the observed change in *pitx2* expression in *pitx2^ΔCE5-7^* was less significant than in the *pitx2^ΔCE5-11^* eyes where a statistically significant decrease was observed at all three stages (2-4-dpf) (Figure 2). Thus, there seems to be a correlation between the size of the deletion (and the number of conserved regulatory elements affected) and the effect on *pitx2* expression as well as the degree of the anterior segment reduction/dysgenesis.

**Figure 2.**
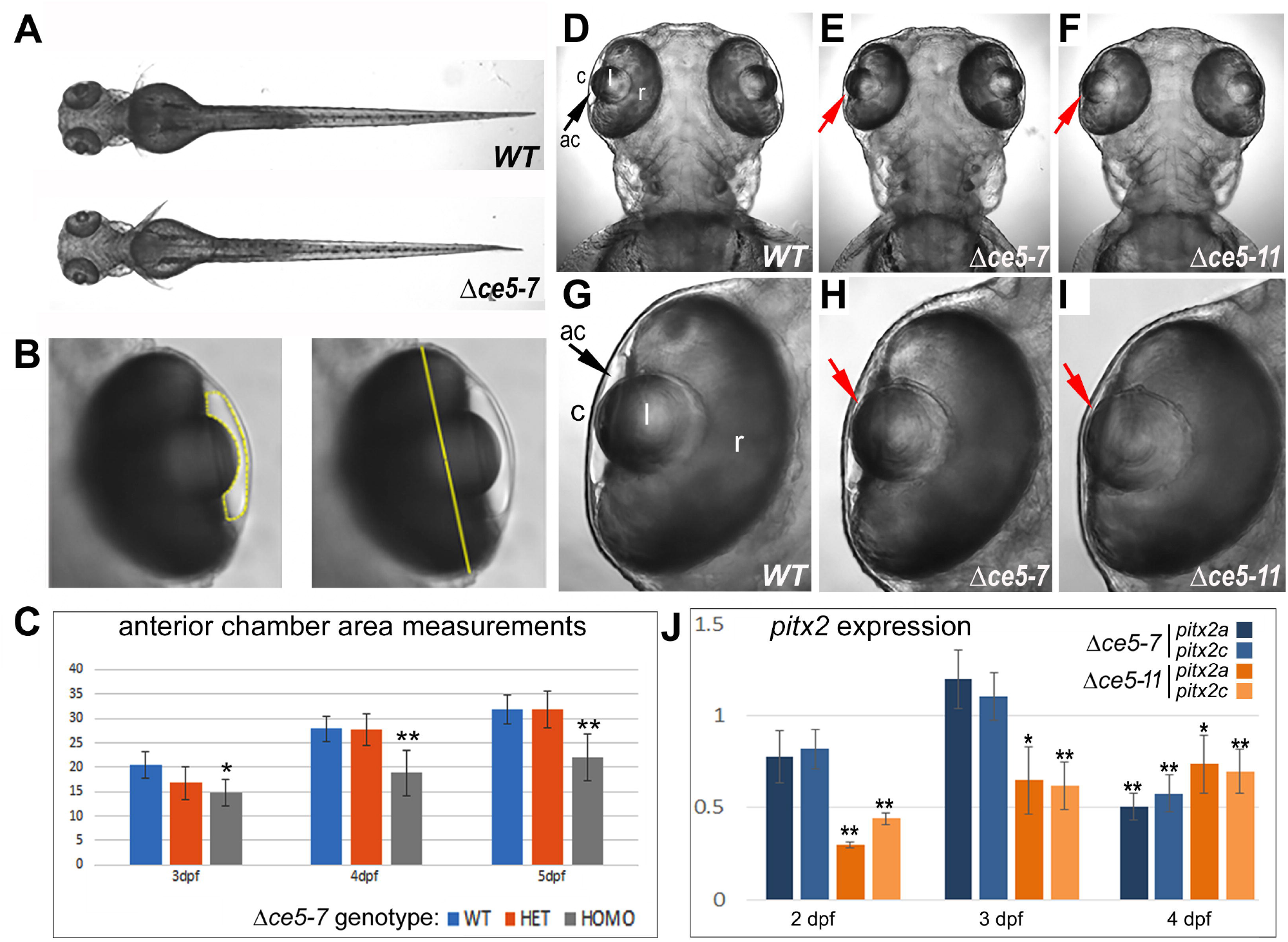
Gross morphology and *pitx2* expression in *pitx2^ΔC5-7^* line. **(A)** Overall growth and morphology of both *pitx2^ΔC5-7^*and *pitx2^ΔC5-11^* mutants at 4-dpf is indistinguishable from WT. **(B)** The anterior chamber area was measured by outlining the interior surface of the cornea, lens and iris using ImageJ (left) and normalizing the value by the width of the eye at the widest point (right). **(C)** Normalized anterior segment measurements show statistically significant differences at 3-dpf (p=0.0019), 4-dpf (0.000044 and 5-dpf (0.000051) between wild-type and homozygous *pitx2^ΔC5-7^* eyes and no statistically significant difference between wild-type and heterozygous eyes for the same mutant (3 dpf-0.019; 4 dpf-0.898; 5dpf-0.985). **(D-l)** Dorsal views of WT and *pitx2^ΔC5-7^* and *pitx2^ΔC5-11^* mutants at 4-dpf show a differing degree of anterior segment space reduction in both mutants. **(J)** qPCR analysis of *pitx2a* and *pitx2c* expression in homozygous eyes of both *pitx2^ΔC5-11^* and *pitx2^ΔC5-7^* mutants compared to WT at 2-, 3- and 4-dpf demonstrate a statistically significant (marked with asterisks) decrease in the expression of *pitx2* transcripts in both mutants at 4-dpf.

### Electron microscopy studies of *pitx2^ΔCE5-7^* and *pitx2^ΔCE5-11^* zebrafish eyes

To obtain an additional insight into the anterior segment phenotype of the deletion mutants, sections of the ventral and dorsal angles of the zebrafish eye as well as the entire cornea were examined by electron microscopy. Wild-type (n=3), homozygous *pitx2^ΔCE5-11^* (n=5) and homozygous *pitx2^ΔCE5-7^* (n=3) 3-dpf mutant eyes as well as wild-type (n=3), homozygous *pitx2^ΔCE5-11^* (n=3) and homozygous *pitx^ΔCE5-7^* (n=3) 14-dpf mutant eyes were analyzed.

At 3-dpf, many structures of the anterior chamber of the zebrafish eye such as the cornea, lens and iris are already developed (Soules and Link 2005). The corneal epithelium at 3-dpf consists of two layers of cells and looks similar in both mutants and WT. The corneal stroma appears to have uniform thickness in both *pitx2^ΔCE5-11^* and *pitx2^ΔCE5-7^* mutants and a continuous endothelial layer is present unlike in the *pitx2^M64*^* mutant, where the endothelial layer is disrupted and corneal stroma is often merged with the lens capsule (Hendee et al. 2018). In the dorsal iridocorneal angle at 3-dpf, the iris stroma is already formed while the non-pigmented epithelial layer is yet to be developed (Soules and Link 2005). The stroma consists of pigmented epithelium containing dark round-shaped melanosomes, iridophores filled with rod-shaped silvery iridosomes, and xanthophores loaded with foam-like pteranosomes. In all WT samples, xanthophores are exterior to the iridophores and are separated from the iridocorneal angle by a layer of undifferentiated cells. In all analyzed *pitx2^ΔCE5-11^* mutant eyes, xanthopohores as well as interspersing undifferentiated cells and occasionally iridophores looked disorganized and were obstructing a significant portion of the dorsal iridocorneal angle. (Figure 3); the dorsal iridocorneal angle in the *pitx2^ΔCE5-7^* mutant also appeared abnormal in 2 out of 3 eyes marked with the presence of large cells of unclear origin next to corneal stroma. These findings are different from the *pitx2^M64*^* homozygous mutant phenotype where the dorsal iris stroma appears to be under-differentiated and the dorsal iridocorneal angle is filled with undifferentiated cells and cellular debris (Hendee at al. 2018). In the ventral iridocorneal angle of WT eyes, differentiating endothelial cells form an orderly single-cell layer next to the corneal stroma. Instead, in the *pitx2^ΔCE5-11^* mutant eyes multi-layered cell sheets underlying the stroma fill up most of the available space in the ventral iridocorneal angle in all examined eyes (Figure 3); the ventral iridocorneal angle of *pitx2^ΔCE5-7^* mutants lacked the layer of undifferentiated cells which typically envelope the tip of the iris stroma (n=2) or had a normal angle appearance (n=1). The undifferentiated mesenchymal cells at the margins of the iridocorneal angles eventually form the endothelium layer of the annular ligament. Similar (while more severe) defects in the ventral angle were present in the in *pitx2^M64*^* mutant (Hendee et al. 2018). The observed features indicate a delay in anterior chamber development, particularly the iridocorneal angles in both mutants.

**Figure 3.**
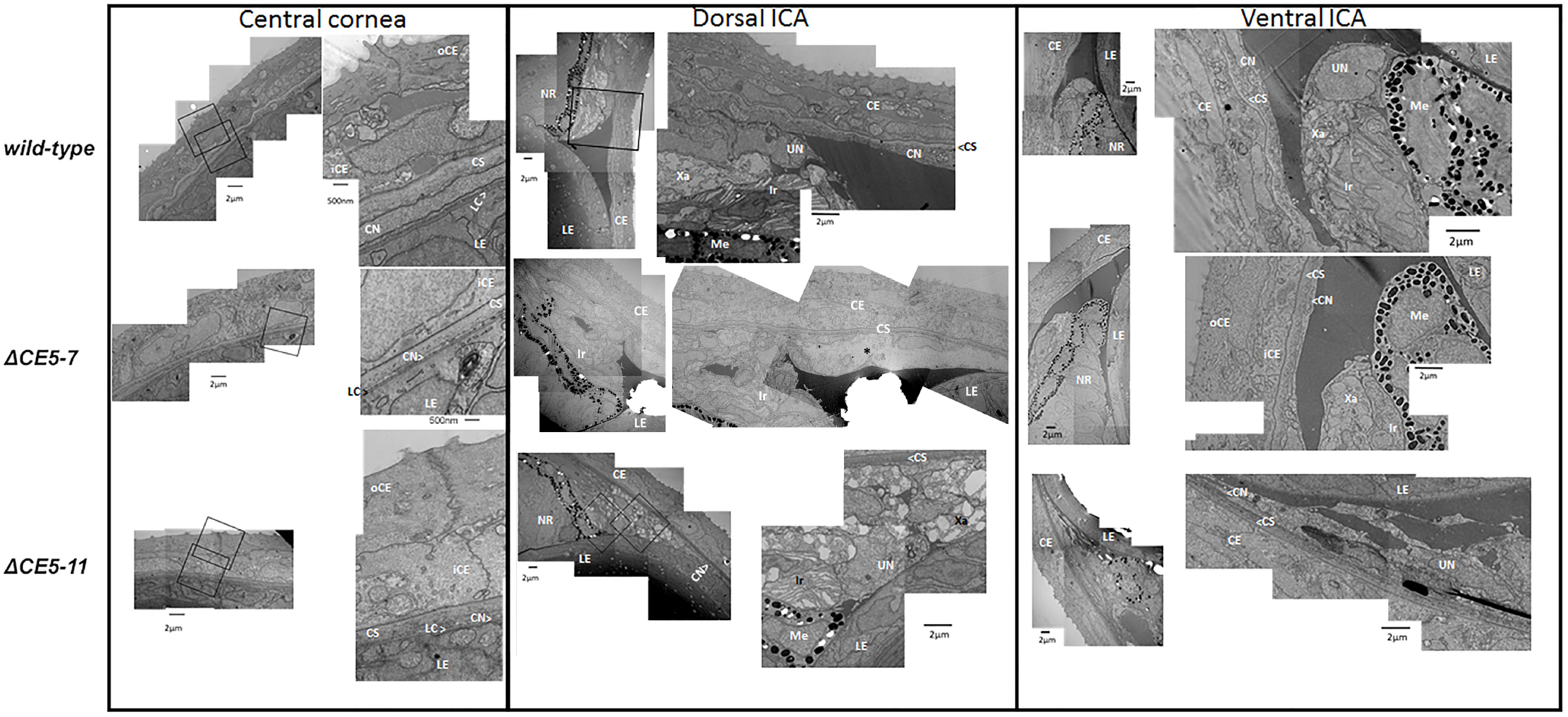
Electron microscopy of *pitx2^ΔC5-7^* and *pitx2^ΔC5-11^* mutant and WT eyes at 3-dpf. Data for the central corneal region as well as both iridocorneal angles (ICA; dorsal and ventral) are shown. Please note abnormalities in the dorsal and ventral angles in both mutants (see text). CE, corneal epithelium; CN, corneal endothelium; CS-corneal stroma; iCE-inner corneal epithelium layer; Ir, iridophore; LC-lens capsule, LE-lens; Me, melanocyte; NR, neural retina; oCE-outer corneal epithelium layer; UN, undifferentiated cell; Xa, xanthophore; * marks cells of unknown origin next to corneal stroma in the dorsal angle of *pitx2^ΔC5-7^* mutant.

At 14-dpf, the WT corneal epithelium is at least 3-cell-layers thick with adjacent cells tightly connected to each other. The *pitx2^ΔCE5-11^* mutant eyes appear to have a thinner corneal epithelial layer with many dead or dying cells in the outer layer (Figure 4); there were some parts of the cornea with only one layer of epithelial cells at its surface. The cornea in the *pitx2*^ΔCE5-7^ mutant looked normal in all samples. The annular ligament, the structural analog of the mammalian trabecular meshwork in in zebrafish, is not fully developed until 17-dpf (Soules and Link 2005). At 14-dpf, mesenchymal cells differentiating into the annual ligament structures are present at the dorsal and ventral angles in the WT and both mutants with no visible differences in their distribution as well as the overall morphology of both angles between WT and mutant larvae (Figure 4).

**Figure 4.**
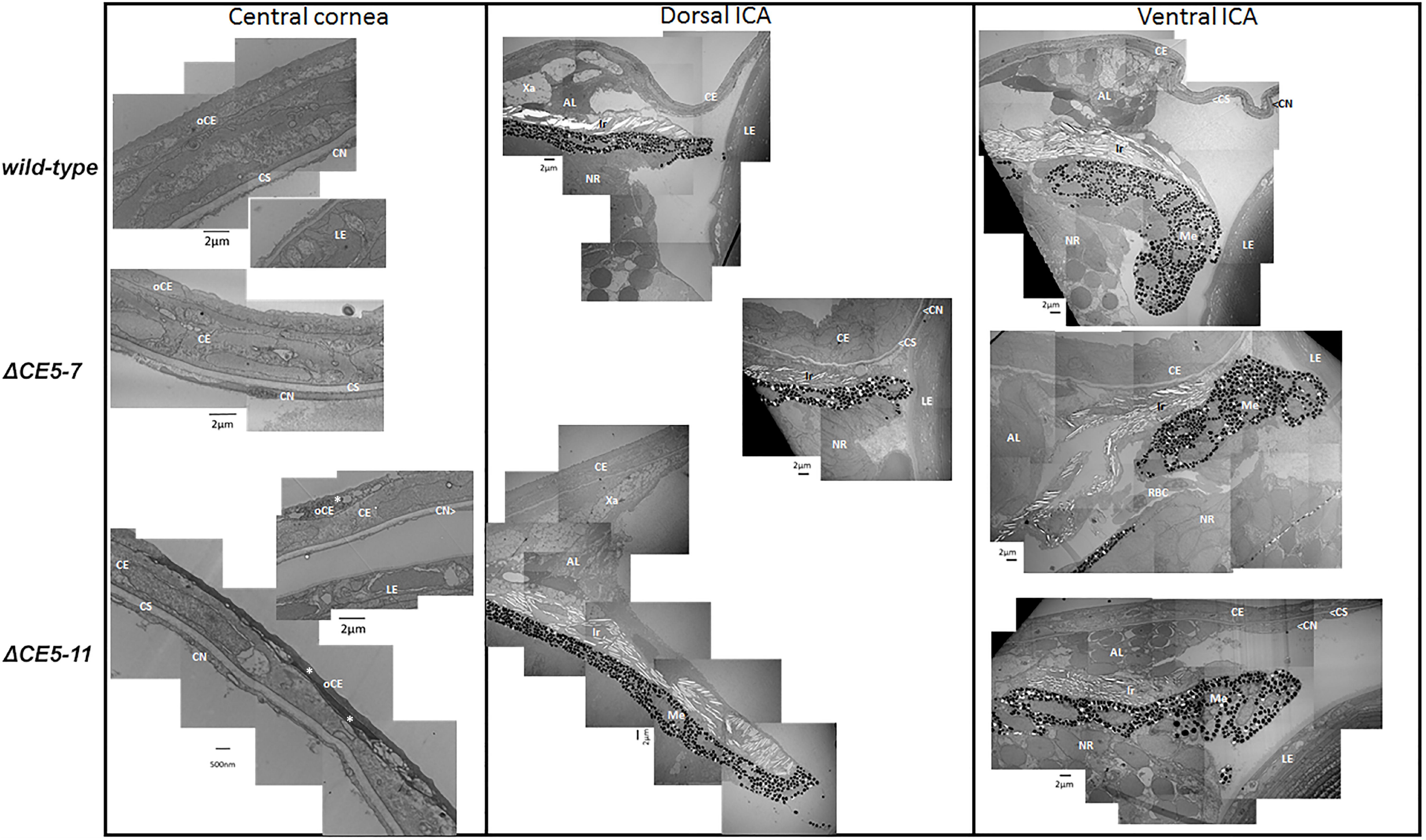
Electron microscopy of *pitx2^ΔC5-7^ and pitx2^ΔC5-11^* mutant and WT eyes at 14-dpf. Data for the central corneal region as well as both iridocorneal angles (ICA; dorsal and ventral) are shown. Both angles appear to have a normal appearance in the mutants; the central cornea shows dying cells (asterisks) in *pitx2^ΔC5-11^* mutant eyes while *pitx2^ΔC5-7^* appears to be normal. AL-differentiating annual ligament cells; CE, corneal epithelium; CN, corneal endothelium; CS-corneal stroma; iCE-inner corneal epithelium layer; LE-lens; Ir, iridophore; Me, melanocyte; NR, neural retina; oCE-outer corneal epithelium layer; UN, undifferentiated cell; Xa, xanthophore.

### Studies of adult mutants and variability of *pitx2^ΔCE5-11^* phenotypes

To investigate the progression of the phenotype, 5-dpf mutant embryos were placed on the system and raised to adulthood. We observed the following survival to adulthood: 31% and 61% for homozygous and heterozygous *pitx2^ΔCE5-11^* siblings, respectively, and 60% for both homozygous and heterozygous *pitx2^ΔCE5-7^* fish; a similar 60-65% survival rate was observed for groups of wild-type embryos. Adult heterozygous or homozygous fish for either deletion appeared to be generally unaffected based on gross observations under a microscope as well as histological studies. However, homozygous males carrying the *pitx2^ΔCE5-11^* allele failed to produce any offspring despite multiple matings, similar to the previously reported *pitx2^M64^\** males (Hendee et al. 2018). The higher rate of larval death among homozygous *pitx2^ΔCE5-11^* mutants is reminiscent of the poor survival noted in *pitx2^M64^\** fish (Hendee et al. 2018) and indicates that partial *pitx2* deficiency (due to the regulatory deletion in *pitx2^ΔCE5-11^*) sensitizes these animals to the effects of additional modifiers that may amplify the negative effects of altered *pitx2* expression.

Consistent with this possibility, we observed a more pronounced ocular phenotype in a subset of *pitx2^ΔCE5-11^* homozygous mutants (Figure 5). Initially, this phenotype was observed in up to 5% of homozygous animals in different breedings, however, the frequency increased up to 25% with select breedings using animals that showed the most significant reduction in the anterior segment as embryos. The more severely affected fish presented with a temporally elongated eye evident starting at 4-6 dpf, ventral iris coloboma in some, extremely narrow anterior segment, and microphthalmia (unilateral in 85% of cases). The more severely affected fish showed a reduced survival, with the majority of animals dying before 1.5 mpf; thus, the surviving adults for either deletion, *pitx2^ΔCE5-7^* mutants or *pitx2^ΔCE5-11^* appear to be grossly normal as noted above. The more severe phenotype observed in a subset of *pitx2^ΔCE5-11^* homozygous shows overlap with the features identified in the *pitx2^M64*^* loss-of-function mutants and, similarly to the above, suggests a sensitizing role for incomplete *pitx2* deficiency and the involvement of additional genetic modifiers which can now be sought in future studies.

**Figure 5.**
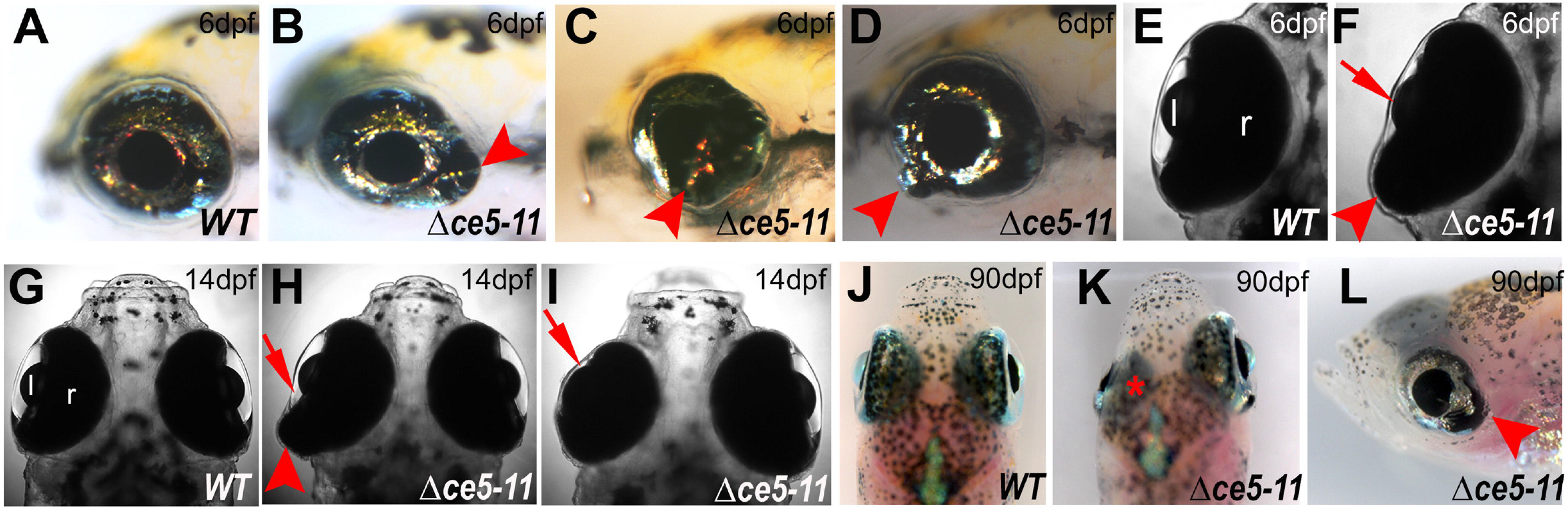
Severe ocular phenotypes observed in a subset of *pitx2^ΔC5-11^* homozygous embryos at 6-dpf **(A-F)**, 14-dpf **(G-l)** and 90-dpf **(J-L)**. Please note temporally elongated retina, coloboma, and other retinal defects (red arrowheads), shallow anterior segment and small eye (red arrows), and mispositioning of the malformed eye (asterisk) similar to the features observed in the *pitx2* loss-of-function line *pitx2^M64*^* (Hendee et al. 2018). The observed severe phenotype was unilateral in the majority (85%) of affected fish.

### DNA methylation analysis of the *pitx2* region in wild-type and *pitx2^ΔCE5-7^* eyes

Bisulfite sequencing was performed using total DNA extracted from 3-dpf *pitx2^ΔCE5-7^* mutant and wild-type eyes with 3 biological replicas for each genotype. For quantification of the methylation level, the cleaned reads were first mapped to the reference Danio rerio GRCz11 genome and then the methylation state of all cytosines in a CpG, CHH, or CHG context was calculated as a ratio of C/C+T for each position for all reads aligned using methyratio.py in BSMAP. The MethylRatio values of all samples were normalized using median scaling normalization.

Methylation status for the *pitx2* region was analyzged by bisulfite sequencing in WT and *pitx2^ΔCE5-7^* 3-dpf eyes. We employed the Integrative Genomics Viewer tool (Thorvaldsdóttir et al. 2013) to visualize the methylation status of all cytosines at the relevant context at single base resolution simultaneously in all samples. Two hypermethylated clusters separated by about 400-bp were mapped to intron 2 in the *pitx2a* transcript ENSDART00000148319.4 in the *pitx2^ΔC5-7^* mutant eyes and no methylation of this region was detected in WT eyes (Figure 6). The first cluster (region 1; Figure 6) overlapped the putative promoter region and 5'-UTR of the *pitx2c* transcript ENSDART00000052569.7. Since hypermethylation of a promoter usually indicates its repression, this result emphasizes the role of the regulatory region within CE5-7 boundaries in expression of *pitx2c* at this stage of embryonic development of the eye. An upstream cluster (region 2; Figure 6) of the hypermethylated CpG in the mutant *pitx2^ΔC5-7^* eyes is situated in the chr 14:36,224,092-36,225,480 interval placing this CpG island outside of the *pitx2c* proximal promoter region; however, hypermethylation of this region in the mutants suggests its importance for expression of one or possibly both *pitx2* isoforms.

**Figure 6.**
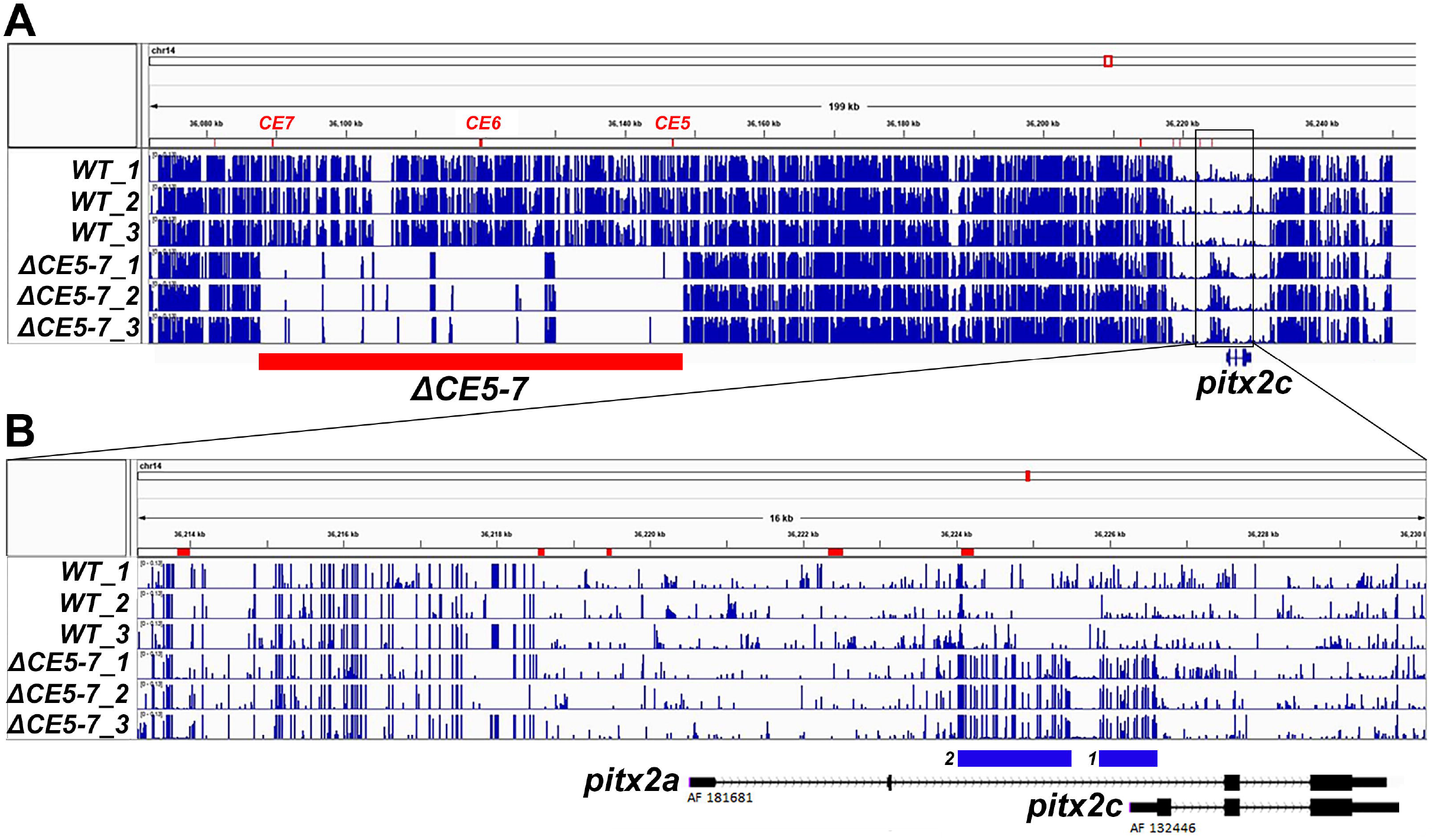
DNA methylation profiles at the *pitx2* region in *pitx2^ΔC5-7^* 3-dpf mutant and WT eyes. **(A)** The DNA methylation status was visualized using Integrative Genomics Viewer tool (Thorvaldsdóttir et al. 2013): three biological repeats for wild-type eyes (top three rows) were compared to three biological repeats for *pitx2^ΔC5-7^* homozygous mutant eyes (bottom three rows) at 3 dpf. Please note CE5-7 deletion region marked with the red bar with all three mutant samples showing no specific reads for this region; at the same time, a region upstream of *pitx2c* (box) displayed hypermethylation in the three mutant samples. **(B)** An enlarged view of this region shows two hypermethylated clusters (numbered blue bars) separated by about 400-bp and located in intron 2 in the *pitx2a* transcript and upstream of *pitx2c.* Cluster 1 overlaps the putative promoter region and 5'-UTR of the *pitx2c* transcript while cluster 2 is located upstream of this region and may be involved in regulation of both isoforms.

## DISCUSSION

The presented data show that deletions of the upstream regulatory elements of *pitx2* are sufficient to induce anterior segment phenotypes in zebrafish, similar to humans (Protas et al. 2017). Consistent with the predicted regulatory roles of the CEs within those regions, these deletions interfere with normal *pitx2* expression levels during embryonic eye development with a positive correlation between the size of the deletion (and number of CEs) and the degree of transcriptional disruption. This corresponds well with the data reported by Volkmann et al. (2011) that suggested that no single CE of *pitx2* could explain the complex endogenous expression pattern seen using *in situ* hybridization in WT embryos. The presented data demonstrated the existence of multiple single elements that could drive expression of GFP in the periocular mesenchyme and when these single elements were combined into the same vector, the number of periocular mesenchymal cells positive for GFP increased. The results of the current study support these observations. We show that *pitx2* expression is decreased when CE5-7 are deleted and is even further reduced when that deletion is extended to CE5-11.

The DNA methylation studies performed in the line carrying the CE5-7 deletion (orthologous to the smallest known human deletion associated with ARS) identified hypermethylation of the *pitx2c* promoter and an additional region upstream of *pitx2c* and internal to *pitx2a* (within intron 2) in embryonic eyes of homozygous mutants at 3-dpf, prior to the significant decrease in the levels of both transcripts identified in 4-dpf mutant eyes. The identified hypermethylation pattern at the *pitx2* promoter and an additional nearby internal regulatory region are likely to explain the decrease in *pitx2* transcript level as strong DNA methylation of promoters is generally associated with transcriptional repression (Miranda and Jones, 2007).

The phenotypes observed in the deletion lines, *pitx2^ΔCE5-7^* and *pitx2^ΔCE5-11^*, included variable reduction of the anterior segment space, with a greater reduction associated with the larger, ΔCE5-11, deletion and a milder phenotype detected in *pitx2^ΔCE5-7^* animals. The observed phenotype is consistent with that seen in embryos with a complete loss of *pitx2* function, *pitx2^M64*^*, with the *pitx2^M64*^* mutants displaying the most severe end of this phenotypic spectrum characterized by the greatest initial anterior segment reduction and a failure to recover from it with highly abnormal anterior segment structures in embryos and all surviving adults (Hendee et al. 2018). The *pitx2^ΔCE5-7^*and *pitx2^ΔCE5-11^* mutants surviving to adulthood (with normal survival in *pitx2^ΔCE5-7^* and reduced rates in *pitx2^ΔCE5-11^* animals) appear normal, including ocular morphology, which suggests that the observed delay/abnormality in their eye development is transient. This may indicate that the *pitx2* deficiency associated with the studied regulatory deletions is transient and becomes resolved at later stages through the involvement of other regulatory regions of *pitx2* or an activation of compensatory pathways leading to normalization of *pitx2* (or its target) expression and recovery from the initial negative impact of *pitx2* downregulation. Additional investigations are likely to provide deeper insight into *pitx2* regulation and interacting pathways.

Interestingly, a subset of *pitx2^ΔCE5-11^* embryos (with enrichment in the progeny of adults that showed the most profound embryonic anterior segment reduction) was found to present with a more severe phenotype characterized by retinal malformations, small eye and a more pronounced/longer lasting reduction of the anterior segment space. These embryos were also characterized by markedly reduced survival. The observed features show similarities to the *pitx2* complete loss of function phenotype and appear to be heritable, suggesting the presence of negative (or absence of positive) modifiers of the pitx2 pathway resulting in a more severe phenotype in these fish. These modifiers can now be identified and may provide insight into the variability of human phenotypes associated with *PITX2* deficiency.

In summary, we report functional evaluation of the smallest regulatory deletion associated with human ARS in zebrafish, ΔCE5-7. The conserved function of this region is highlighted the presence of the anterior segment phenotype in homozygous zebrafish mutants, similar to the previously reported phenotypes associated with a larger, ΔCE5-11, deletion. Identifying the factors which bind to the conserved elements or other sequences contained within those regions is likely to inform our understanding of eye development and potentially reveal novel genes implicated in ASD. Additionally, the observed phenotypic variability associated with *pitx2* deficiency due to regulatory mutations may assist in the identification of factors that modify the effects of these mutations.

## ACKNOWLEDGEMENTS

This work was supported by NEI grants R01EY025718, R01EY015518 (EVS) as well as funds provided by the Children's Research Institute Foundation at Children's Hospital of Wisconsin (EVS). The authors would like to thank Linda M. Reis, MS, CGC, for a careful reading and editing of the manuscript and Samuel Thompson, BS, for his help with zebrafish genotyping.

## SUPPLEMENTAL DATA

**Supplemental Table 1**. Reagents utilized to generate and to genotype zebrafish carrying CE5-7 deletion.

