## Supplemental Table 1 for "Deletions of distant regulatory sequences upstream of zebrafish *pitx2* result in a range of ocular phenotypes"

**Table S1.**

| **sgRNA** | **Target Site** | **Oligo 1** | **Oligo 2** |
| --- | --- | --- | --- |
| CE5 Downstream | GGACACGCTGAAGAAAAGCT | TAGGACACGCTGAAGAAAAGCT | AAACAGCTTTTCTTCAGCGTGT |
| CE7 Upstream | GGGATTCAGTATCGTATCCC | TAGGGATTCAGTATCGTATCCC | AAACGGGATACGATACTGAATC |
| ***pitx2 CE 5-7* Deletion Line Primers:** |  |  |  |
| A combination of 1803 and 1713 identifies Wildtype Allele | | | |
| A combination of 1712 and 1713 identifies CE5-7 Deletion Allele | | | |
| 1803 | CE8-11_del_R | TTTGGGGGTGGGAAAAGGAC |  |
| 1712 | CE5_sg2_screen_R | GCGCTTTGCTGCAGATTGTA |  |
| 1713 | CE7_sg1_screen_F | GCGAACAAAGAGCTCAGCAC |  |
